# Androgen receptors expressed in the primary sensory neurons regulate mechanical pain sensitivity

**DOI:** 10.1101/2025.03.13.642983

**Authors:** Fumihiro Saika, Daisuke Uta, Yohji Fukazawa, Yuko Hino, Yu Hatano, Shiroh Kishioka, Hiroyuki Nawa, Shinjiro Hino, Kentaro Suzuki, Norikazu Kiguchi

## Abstract

The expression of hormonal receptors in pain-processing regions complicates understanding the hormonal effects on pain mechanisms. This study investigates androgen receptor (AR) involvement in pain sensitivity and sex differences in pain perception. Mechanical pain thresholds were higher in normal male mice compared to gonadectomized (GDX) male and normal female mice, correlating with serum testosterone levels. In the dorsal root ganglia (DRG), AR was expressed in normal males but undetectable in GDX males and normal females. In male sensory neuron-selective AR conditional knockout (AR-cKO) mice, mechanical pain thresholds were significantly lower than in wild-type males. In female mice, administration of testosterone propionate or dihydrotestosterone significantly raised mechanical pain thresholds, accompanied by increased AR expression in the DRG. This effect was abolished in AR-cKO females, consistent with male findings. These results indicate that primary sensory neurons are critical targets of androgen signaling in regulating mechanical pain sensitivity.

## Introduction

Pain is a critical alert system embedded in both humans and animals, essential for recognizing potential tissue damage or injury, and triggering protective responses ^1,2^. While it was traditionally assumed that pain mechanisms were similar across sexes, accumulating evidence indicates significant sex-based differences in pain perception and tolerance in both preclinical research and clinical practice ^3^. Females typically exhibit lower pain thresholds and tolerance in experimental settings, reflecting greater sensitivity to painful stimuli ^4–7^. These sex differences are also observed in chronic pain conditions like fibromyalgia and migraines, which are more prevalent and severe in females compared to males ^4,8^. Despite multiple studies on pain perception, it remains unclear which types of painful stimuli consistently show sex differences under physiological conditions. Moreover, the mechanisms underlying these differences are still under investigation, likely involving complex interactions among sex hormones, sex-specific genes, and neural networks and activities ^3,8,9^. Of particular interest are the roles of sex hormones (e.g., androgens and estrogens) in pain perception, a topic gaining increased attention in pain research ^10,11^.

Several lines of evidence suggest that androgens and estrogens modulate pain perception ^9,12,13^. Due to the widespread expression of their receptors and biosynthesis process in which androgens are converted to estrogens ^14–16^, understanding their hormonal effects is complex. We previously reported that pexidartinib exhibited dimorphic therapeutic effects on chronic pain between normal male and gonadectomized (GDX) male mice ^17^, suggesting androgens play a key role in sex differences in pain. However, the molecular mechanisms by which androgens regulate the pain sensitivity remain unclear.

The androgen receptor (AR) is widely expressed in regions involved in pain transmission and processing, including the brain, spinal cord, and dorsal root ganglia (DRG) ^15,18^. Since physiological pain often originates from the activation of primary sensory neurons and is transmitted to the central nervous system (CNS) ^19,20^, it is crucial to explore the androgen targets within the pain transmission pathway. As primary sensory neurons detect the type and intensity of pain ^20,21^, we hypothesized that androgens regulate sensory neuron characteristics and contribute to sex differences in pain perception.

To test this hypothesis, we examined the relationship between androgen levels and pain responses to various painful stimuli using GDX male mice, which lack circulating androgens, and female mice treated with androgens. To determine whether androgens directly affect primary sensory neurons, we generated sensory neuron-selective AR conditional knockout (AR-cKO) male and female mice. This approach allows us to isolate the effects of androgens on sensory neurons from those on CNS neurons. Importantly, AR-cKO mice serve as a precise tool for evaluating the role of androgens in pain regulation while excluding the potential effects of estrogens converted from androgens.

## Results

### Mechanical pain thresholds in a sex- and androgen-dependent manner

We demonstrated sex- and androgen-dependent differences in pain responses across various modalities, including mechanical, thermal, and chemical stimuli. The mechanical pain threshold assessed using the von Frey test was significantly higher in male mice compared to female mice. To investigate the relationship between androgen levels and mechanical pain thresholds, we evaluated bilaterally gonadectomized (complete GDX) and unilaterally gonadectomized (partial GDX) male mice four weeks post-surgery. Partial GDX males exhibited a slight decrease in mechanical pain threshold, while complete GDX males showed a marked reduction, reaching levels comparable to those of females (**Fig. 1a**). Serum testosterone levels were highest in normal males, followed by partial GDX males and lowest in complete GDX males. In contrast, females had significantly lower testosterone levels than normal males (**Fig. 1b**). There were no significant differences in thermal pain latency, as evaluated by the Hargreaves test (**Fig. 1c**), nor in chemical pain-related behavior during both the first (0–10 min) and second (10–50 min) phases induced by formalin injection (**Fig. 1d**) between normal males and females. These results suggest that mechanical pain thresholds, but not thermal or chemical pain modalities, correlate with serum testosterone levels, indicating that androgens play a regulatory role in mechanical pain sensitivity.

**Fig. 1.**
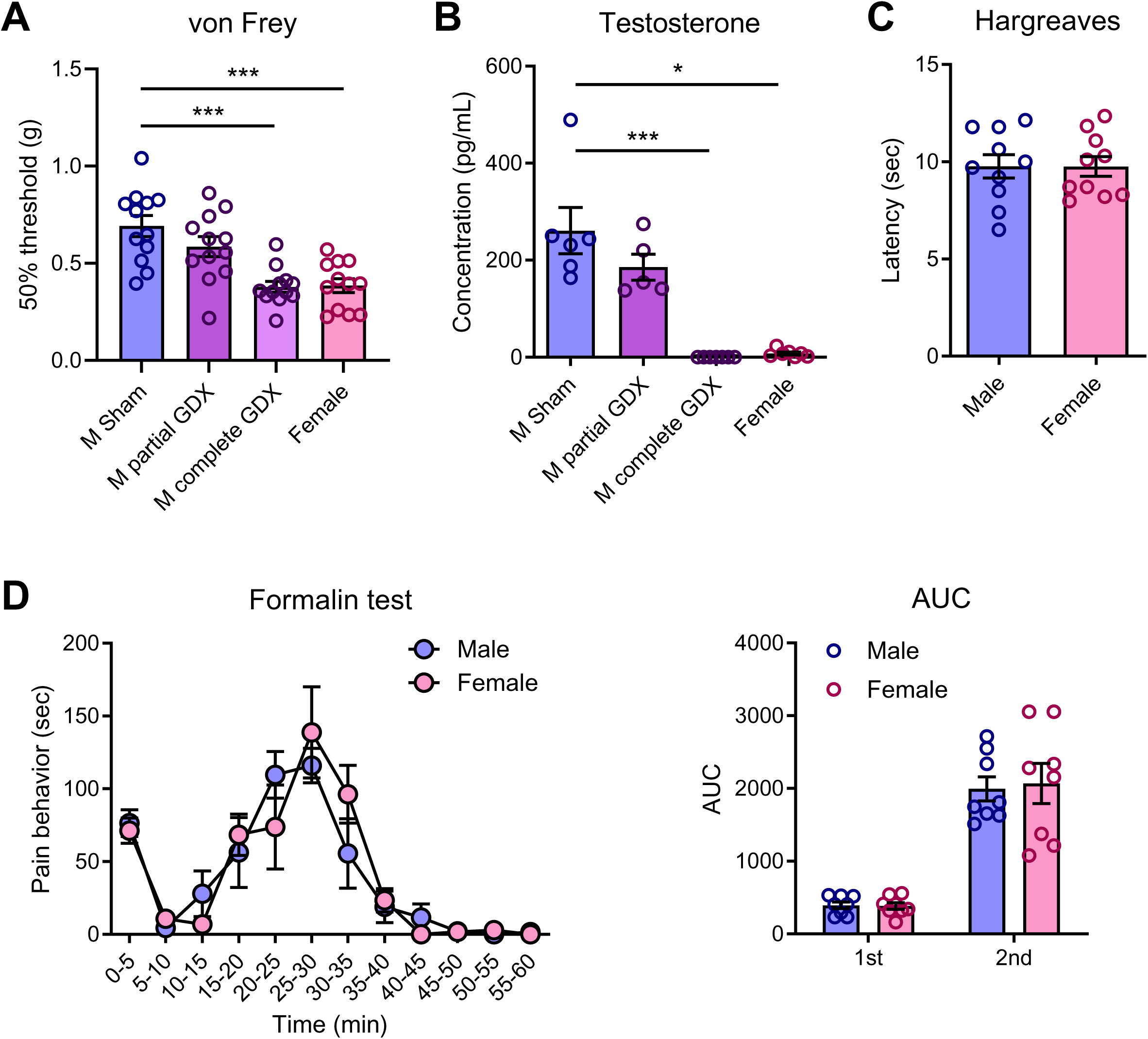
Changes in the mechanical pain threshold in a sex- and androgen-dependent manner. Both male and female mice were used, with male mice undergoing gonadectomy (GDX). (a) Mechanical pain thresholds in normal male, partial GDX male, complete GDX male (4 weeks post-surgery), and female mice were assessed using the up-down method with the von Frey test (n=12 mice). Statistical analysis was conducted using one-way analysis of variance (ANOVA), followed by Dunnett’s multiple comparison test (***P<0.001). (b) Serum testosterone levels were measured using LC–MS/MS (n=5–7 mice, Kruskal-Wallis’s test followed by Dunn’s multiple comparison test, ***P<0.001, *P=0.0322). Thermal pain latency in male and female mice was evaluated using the Hargreaves test (n=10 mice, Student’s t-test). (d) Chemical pain responses in male and female mice after intraplantar injection of formalin. (n=8 mice, Mann-Whitney’s U test).

### Androgen levels and the excitability of pain-responsive neurons

We investigated whether androgen levels influence the excitability of pain-processing neurons in the superficial layer of the spinal dorsal horn (SDH) using electrophysiological techniques. The spontaneous firing rates of SDH neurons were similar between normal males and females (**Fig. 2a**). However, the spontaneous firing rates in GDX males were significantly higher compared to normal males (**Fig. 2b**). The firing rates of SDH neurons in response to 0.16 g and 0.4 g mechanical stimuli were significantly greater in females compared to normal males (**Fig. 2c**). Additionally, firing rates of SDH neurons in response to all tested filament forces were elevated in GDX males compared to normal males (**Fig. 2d**). These results suggest that androgen levels are associated with the excitability of mechanical pain-responsive SDH neurons.

**Fig. 2.**
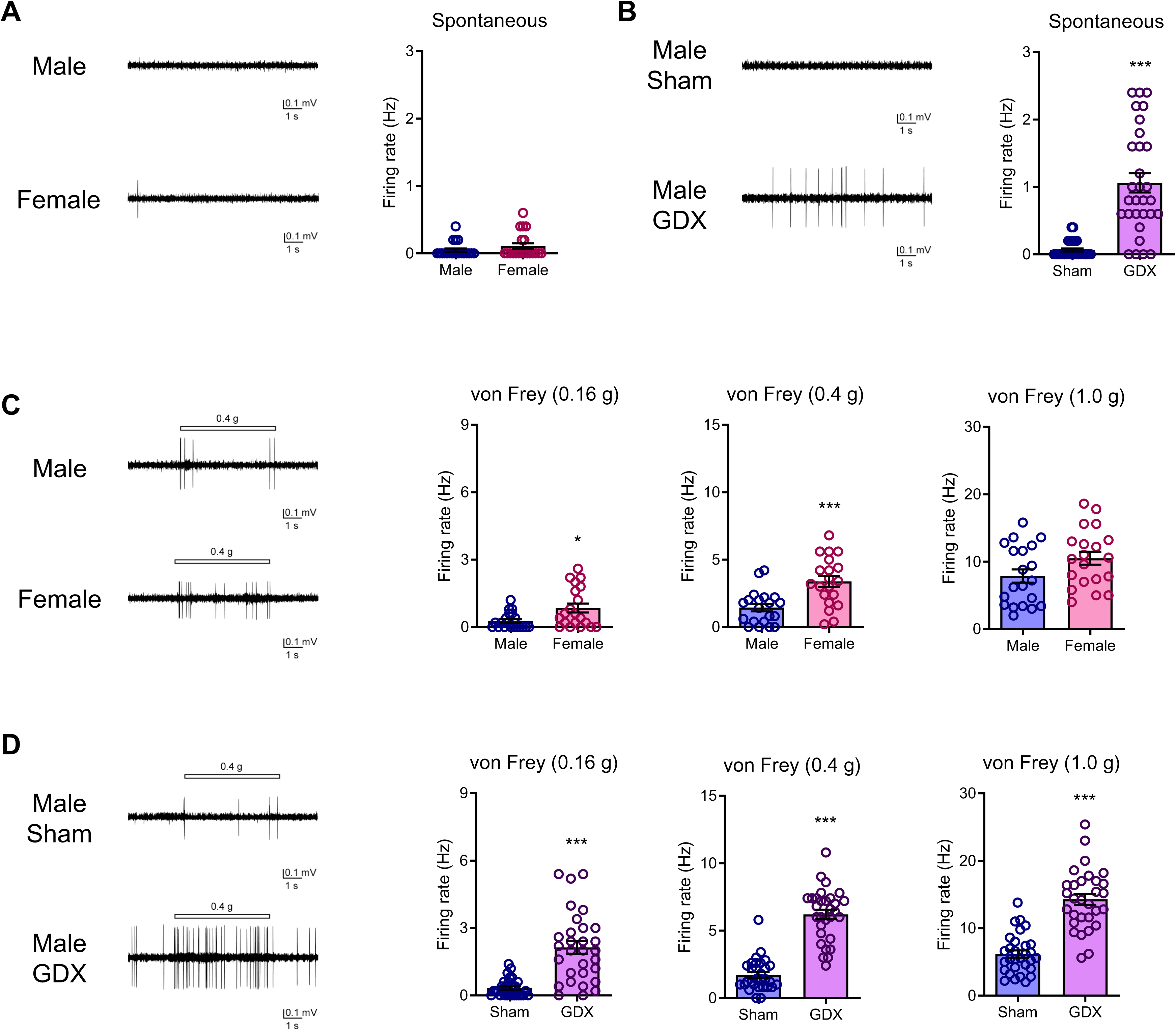
Correlation between androgen levels and the excitability of pain-responsive spinal dorsal horn (SDH) neurons. Spontaneous firing and von Frey filament-evoked firing (0.16 g, 0.4 g, 1.0 g) from SDH neurons were recorded *in vivo* using extracellular techniques in normal male, female, and GDX male mice. (a, b) Representative traces and the average spontaneous firing rates from SDH neurons in male and female mice (a, n=20 cells, Mann-Whitney’s U test) or in normal male and GDX male mice (b, n=30 cells, Mann-Whitney’s U test, ***P<0.001). (c, d) Representative traces and the average von Frey filament-evoked firing rates from SDH neurons in male and female mice (c, n=20 cells, Mann-Whitney’s U test, ***P<0.001, *P=0.0101) and in normal male and GDX male mice (d, n=30 cells, Mann-Whitney’s U test, ***P<0.001).

### AR signaling in sensory neurons in a sex- and androgen-dependent manner

DRG neurons include various subsets, such as C-, Aδ-, and Aβ-fibers, which transmit different types of painful stimuli. To identify the primary cellular targets of androgens among these sensory neurons, we examined the distribution of AR in the DRG. AR protein expression was detected in normal males but was undetectable in GDX males and normal females, and it colocalized with NeuN, a neuronal nuclei marker (**Fig. 3a**). Nuclear translocation of the AR suggests activation of androgen signaling. Quantitative analysis revealed that the number of AR^+^ cells in normal males was significantly higher than in GDX males or normal females (**Fig. 3b**). Immunohistochemical analysis showed that AR^+^ neurons were distributed across several neuronal subtypes with varying cell diameters (10–50 μm), including small C-fibers, medium Aδ-fibers, and large Aβ-fibers, with 47.7% (452/948) of DRG neurons expressing AR (**Fig. 3c**). Most AR^+^ cells were also NeuN^+^ (96.2%), and 44.1%, 46.9%, and 8.2% of AR^+^ cells colocalized with neurofilament 200kD (NF200; a myelinated A-fiber marker), calcitonin gene-related peptide (CGRP; a peptidergic C-fiber marker), and Isolectin B4 (IB4; a non-peptidergic C-fiber marker), respectively (**Fig. 3d,e**). Notably, the majority of AR^+^ C-fibers were CGRP^+^ peptidergic subsets compared to IB4-labeled non-peptidergic C-fibers. These findings suggest that AR signaling may modulate nociceptive sensory processing in primary sensory neurons.

**Fig. 3.**
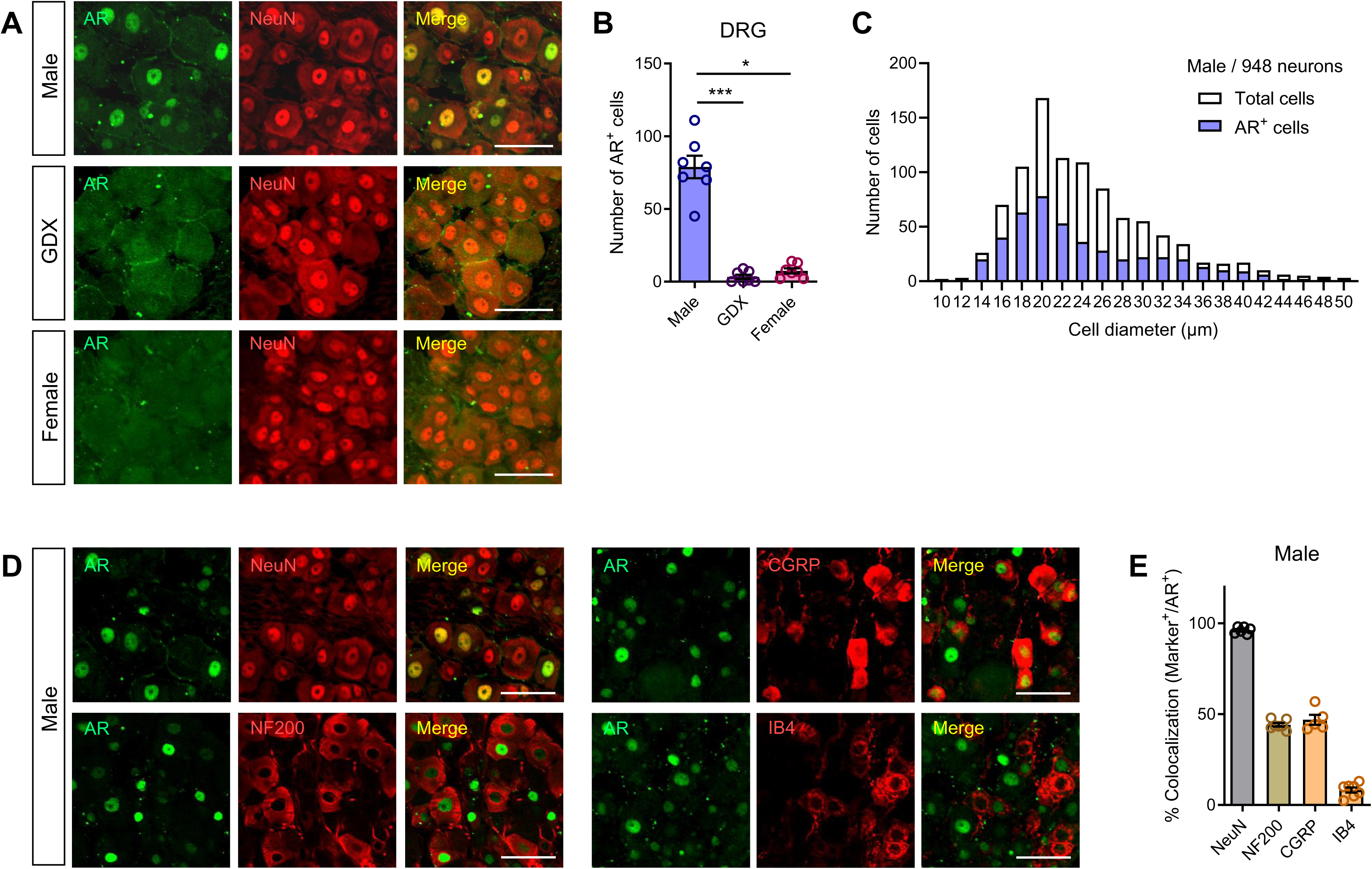
Expression of androgen receptor (AR) in the nuclei of primary sensory neurons in males. (a) Immunohistochemical visualization of AR expression in the dorsal root ganglia (DRG) of normal male, GDX male, and normal female mice was visualized through immunohistochemistry (IHC). (b) Quantitative analysis of the number of AR^+^ cells (n=7 mice, Kruskal-Wallis’s test followed by Dunn’s multiple comparison test, ***P<0.001, *P=0.0192). (c) Distribution rate of AR within 948 DRG neurons of varying cell diameters in male mice. (d) Double immunostaining of AR with neuronal markers (NeuN, NF200, CGRP, or IB4-labeled non-peptidergic C-fibers) in the DRG in male mice. Scale bars = 50 μm (a,d). (e) Quantitative analysis of the percentage of neuronal marker^+^ cells colocalized with AR^+^ cells in the DRG of male mice (n=5–7).

### Roles of AR in sensory neurons for regulating mechanical pain sensitivity

Since Nav1.8 (*Scn10a*) is predominantly expressed in the DRG (**Fig. S1**) and C-fibers ^21,22^, we used Nav1.8-Cre mice ^23^ to perform sensory neuron-selective depletion of AR. In both male and female Nav1.8-Cre::R26-LSL-tdTomato mice, Cre-dependent tdTomato expression was observed in the cell bodies (DRG) and nerve terminals (superficial area of the SDH) of sensory neurons (**Fig. S2a**). Additionally, tdTomato expression partially colocalized with AR in the DRG of males (**Fig. S2b**). Most tdTomato^+^ cells were NeuN^+^, and the colocalization ratio of tdTomato expression in CGRP^+^ or IB4-labeled C-fibers was higher than in NF200-labeled A-fibers (**Fig. S2c**).

As the *AR* gene is located on the X chromosome^24^, Nav1.8-Cre::AR^flox/y^ mice were used as AR-cKO males (**Fig. 4a**). Similar to GDX males and normal females, the mechanical pain threshold in AR-cKO males was significantly lower than in wild-type (WT; AR^flox/y^) males (**Fig. 4b**), while Nav1.8-Cre males exhibited normal pain thresholds (**Fig.S3**). Additionally, no significant differences were observed in thermal pain latency (**Fig. 4c**), chemical pain-related behaviors induced by formalin injection (**Fig. 4d**), nor time spent on the rotarod (**Fig. S4**), indicating that AR-cKO males had pain sensitivity similar to normal females. Immunohistochemical analysis showed a significant reduction in the number of AR^+^ cells in A- and C-fibers in AR-cKO males compared to WT males. The most substantial AR depletion was observed in CGRP^+^ C-fibers, compared to NF200^+^ A-fibers and IB4-labeled C-fibers (**Fig. 4e,f**). These findings suggest that CGRP^+^ C-fibers may be the primary targets of androgens in regulating mechanical pain sensitivity.

**Fig. 4.**
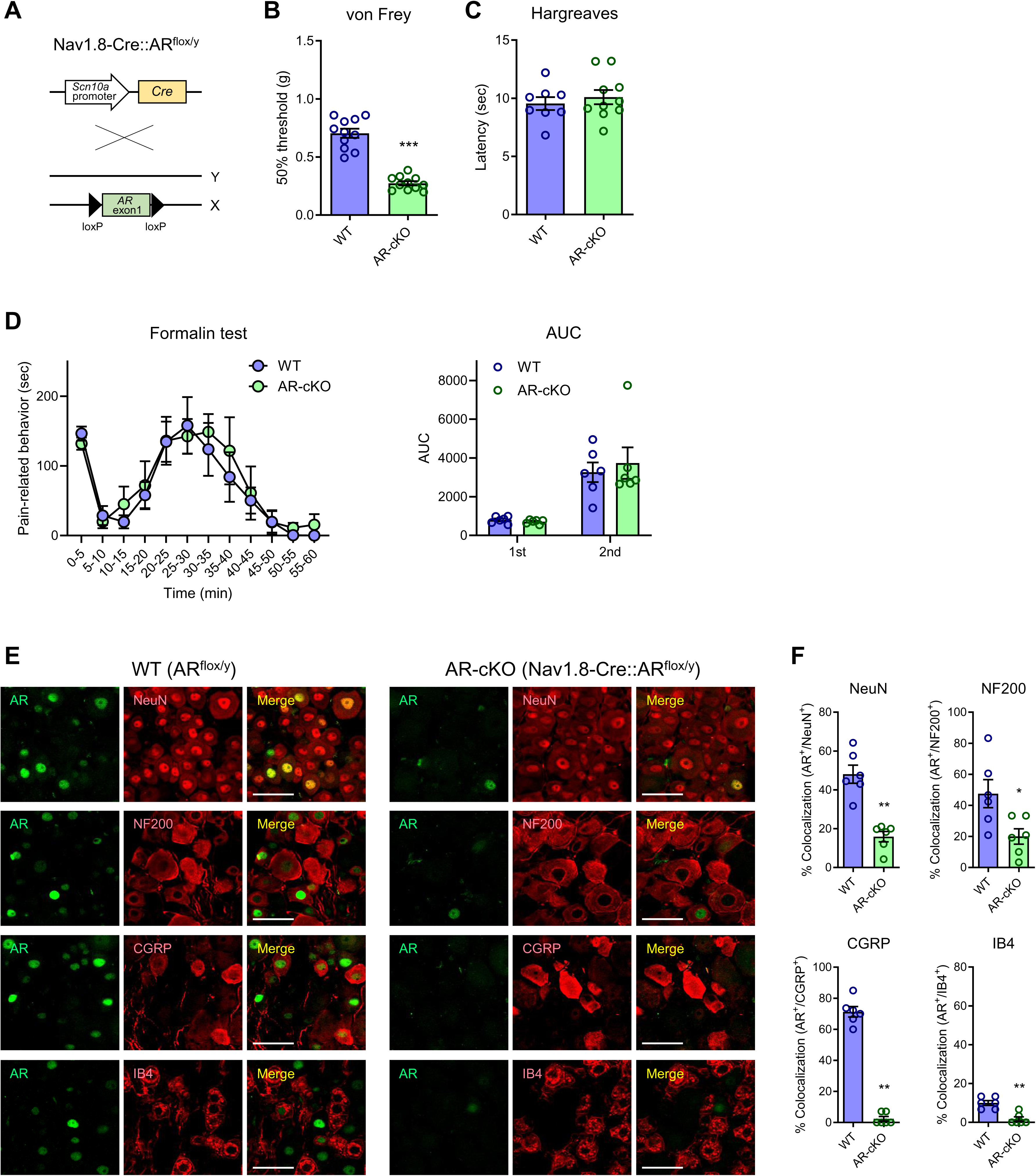
Roles of AR in sensory neurons for regulating mechanical pain sensitivity in males. (a) Nav1.8-Cre mice were crossed with AR^flox/flox^ mice to generate sensory neuron-selective AR conditional knockout (Nav1.8-Cre::AR^flox/y^, AR-cKO) mice. (b) Mechanical pain thresholds in male wild-type (WT) and AR-cKO mice were assessed using the up-down method with the von Frey test (n=11, Welch’s t-test, ***P<0.001). (c) Thermal pain latency in male WT and AR-cKO mice were evaluated using the Hargreaves test (n=8-10 mice, Student’s t-test). Chemical pain responses in male WT and AR-cKO mice were assessed after intraplantar injection of formalin (n=6 mice, Mann-Whitney’s U test). (e) Double immunostaining of AR with neuronal markers (NeuN, NF200, CGRP, or IB4-labeled non-peptidergic C-fibers) in the DRG of male WT and AR-cKO mice. Scale bars = 50 μm. (f) Quantitative analysis of the percentage of AR^+^ cells colocalized with neuronal marker^+^ cells in the DRG of male WT and AR-cKO mice (n=6 mice, Mann-Whitney’s U test, **P=0.0022 (NeuN), *P<0.0325 (NF200), **P=0.0022 (CGRP), **P=0.0065 (IB4)).

### AR depletion in sensory neurons and the excitability of SDH neurons

We sought to obtain electrophysiological evidence that AR in sensory neurons modulates the excitability of pain-responsive neurons in the SDH of AR-cKO mice. The spontaneous firing rates of SDH neurons in AR-cKO males were significantly higher compared to WT males (**Fig. 5a**). Additionally, the firing rates of SDH neurons in response to mechanical pain stimuli were markedly increased at all tested filament forces (0.16, 0.4 and 1.0 g) in AR-cKO males compared to WT males (**Fig. 5b**). These results confirm that AR signaling in primary sensory neurons positively regulates the excitability of mechanical pain-responsive SDH neurons.

**Fig. 5.**
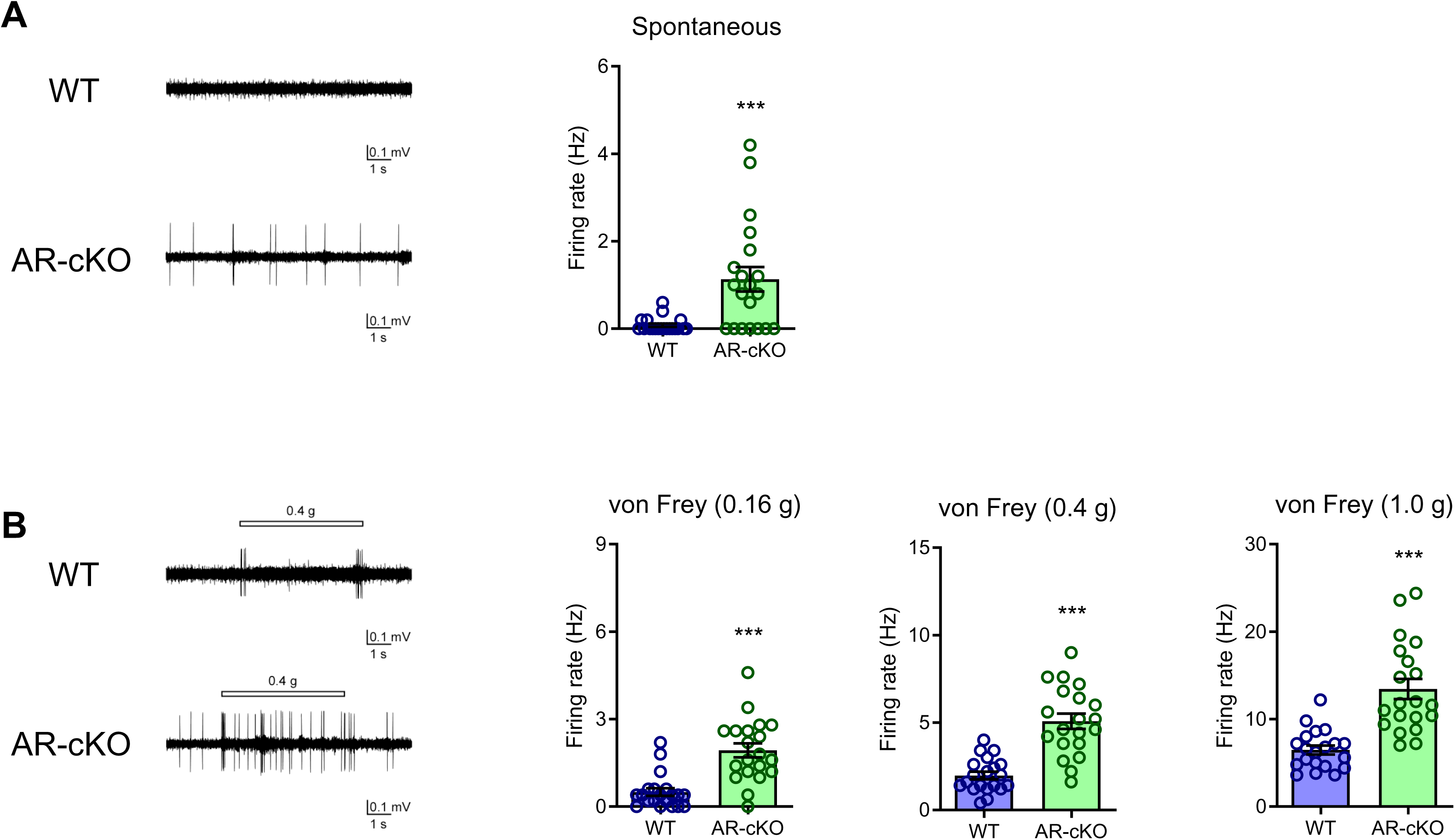
Correlation of AR depletion in sensory neurons and the excitability of pain-responsive SDH neurons. Spontaneous firing and von Frey filament-evoked firing (0.16 g, 0.4 g, 1.0 g) from SDH neurons were recorded *in vivo* using extracellular techniques in male WT and AR-cKO mice. (a) Representative traces and the average spontaneous firing rates from SDH neurons in male WT and AR-cKO mice (n=20 cells, Mann-Whitney’s U test, ***P<0.001). (b) Representative traces and the average von Frey filament-evoked firing rates from SDH neurons in male WT and AR-cKO mice (n=20 cells, Mann-Whitney’s U test, ***P<0.001).

### Exogenous androgen and mechanical pain sensitivity in females

We next investigated the effects of exogenous androgen administration on mechanical pain thresholds in females, which typically exhibit lower endogenous androgen levels. A single systemic intraperitoneal (i.p.) administration of testosterone propionate (TP; 30 or 100 mg/kg) or dihydrotestosterone (DHT; 10, 30 or 100 mg/kg) significantly increased the mechanical pain thresholds on day 1 in a dose-dependent manner, though these effects dissipated by day 2 (**Fig. 6a**). In contrast, DHT administration (100 mg/kg, i.p.) did not affect the threshold in males (**Fig. 6b**), suggesting that the endogenous androgen levels in males are sufficient for regulating of the pain thresholds. Immunohistochemical analysis revealed a marked increase in AR protein expression colocalized with NeuN in the DRG of females on day 1 after DHT administration (100 mg/kg, i.p.) (**Fig. 6c,d**). AR expression emerged and was distributed across several neuronal subtypes with varying cell diameters, and 48.0% (333/694) of DRG neurons expressed AR (**Fig. 6e**). Consistent with observations in males, most AR^+^ cells were NeuN^+^ (97.6%), and 37.8%, 43.7%, and 6.4% of AR^+^ cells colocalized with NF200, CGRP, and IB4, respectively (**Fig. 6f,g**). These results suggest that exogenous androgens can regulate mechanical pain sensitivity in females.

**Fig. 6.**
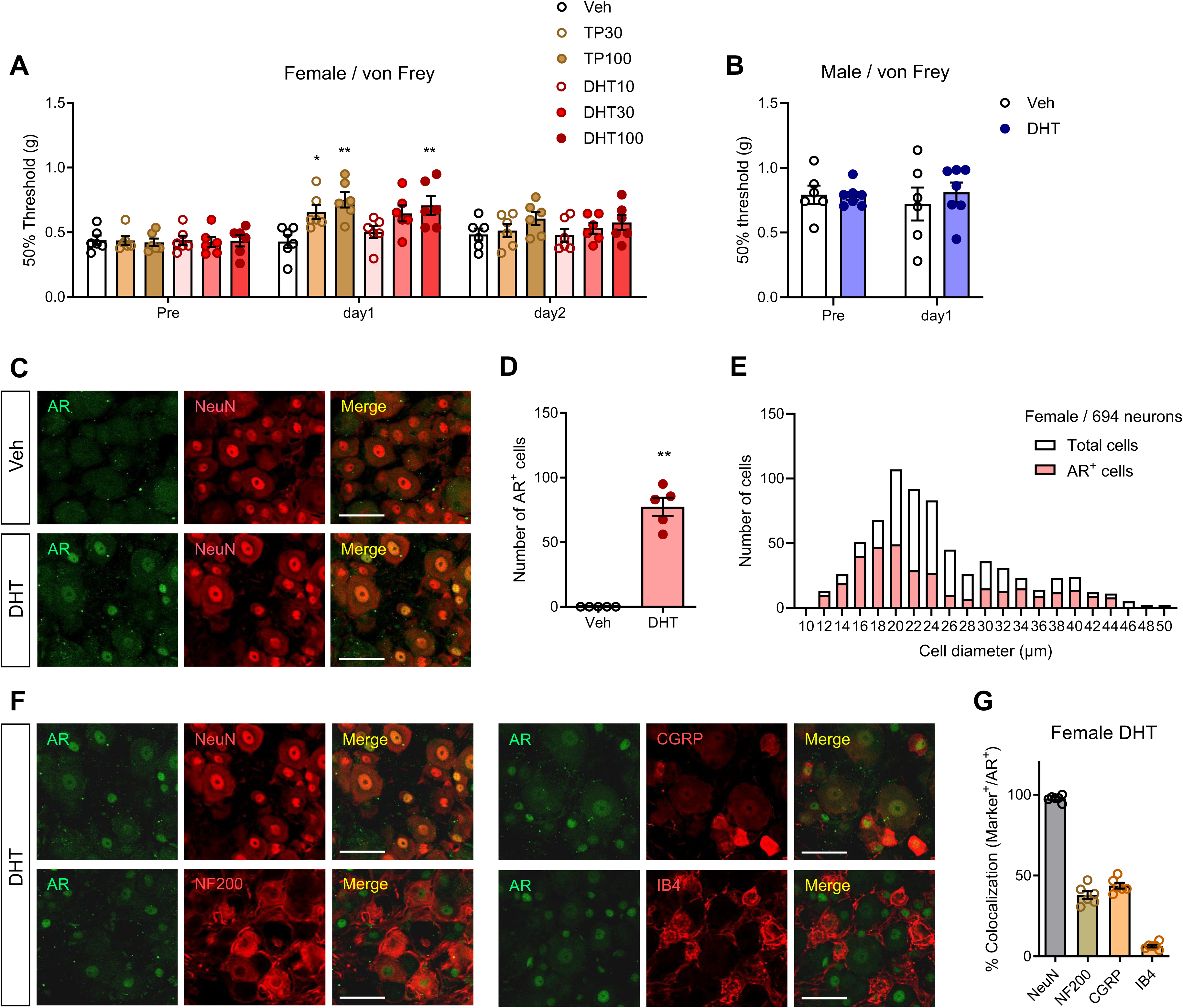
Effects of exogenous androgen administration on mechanical pain sensitivity in females. (a) Mechanical pain threshold in female mice following intraperitoneal (i.p.) administration of testosterone propionate (TP; 30, 100 mg/kg) or dihydrotestosterone (DHT; 10, 30, 100 mg/kg) were assessed using the up-down method with the von Frey test (n=6, one-way ANOVA followed by Dunnett’s multiple comparison test, *P=0.0363, **P=0.002 (TP100), **P=0.0081 (DHT100)). (b) Mechanical pain thresholds in male mice following i.p. administration of DHT (100 mg/kg) were assessed using the up-down method with the von Frey test (n=6-7, Welch’s t-test). (c–g) AR expression in the DRG of female mice on day1 after i.p. administration of DHT were visualized by IHC (c). (d) Quantitative analysis of the number of AR^+^ cells (n=5, Mann-Whitney’s U test, **P=0.0079). (e) Distribution rate of AR within 694 DRG neurons of varying cell diameters. (f) Double immunostaining of AR and neuronal markers (NeuN, NF200, CGRP, or IB4-labeled non-peptidergic C-fibers) in the DRG of female mice on day1 after i.p. administration of DHT. Scale bars = 50 μm (c, f). (g) Quantitative analysis of the percentage of neuronal marker^+^ cells colocalized with AR^+^ cells in the DRG of female mice (n=6).

### Roles of AR in regulating mechanical pain sensitivity in females

To analyze the target cells underlying the pain-regulating effects of DHT in females, we used female sensory neuron-selective AR-cKO mice (Nav1.8-Cre::AR^flox/flox^) (**Fig. 7a**). The increase in mechanical pain threshold on day 1 after DHT administration (100 mg/kg, i.p.) was almost completely abolished in AR-cKO females (**Fig. 7b**), suggesting that AR activation in sensory neurons by exogenous androgens reduces mechanical pain sensitivity. After DHT administration, the number of AR^+^ cells in A- and C-fibers was significantly lower in AR-cKO females compared to WT females, with the most substantial AR depletion in CGRP^+^ neurons (**Fig. 7c,d**). Additionally, we used SDH neuron-selective Cre-expressing (Lbx1-Cre) mice ^25,26^. In Lbx1-Cre::R26-LSL-tdTomato females (**Fig. S5a**), Cre-dependent tdTomato expression was observed in the superficial area of the SDH (**Fig. S5b**). The effects of DHT (100 mg/kg, i.p.) were similar in both WT and SDH neuron-selective AR-cKO females (Lbx1-Cre::AR^flox/flox^) (**Fig. S5c,d**). These findings suggest that AR activation in primary sensory neurons directly regulates mechanical pain sensitivity in both males and females.

**Fig. 7.**
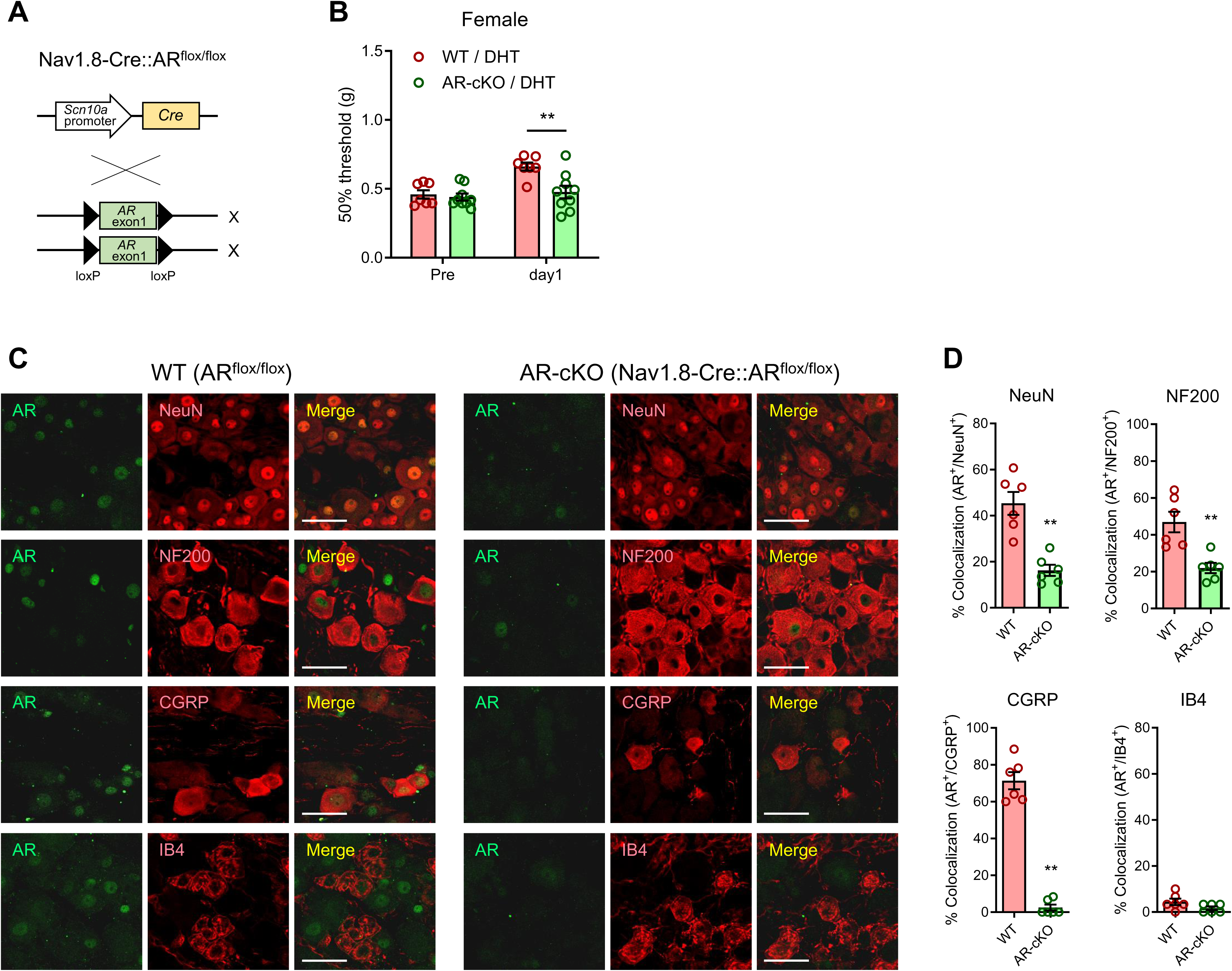
Roles of AR in regulating mechanical pain sensitivity in females. (a) Nav1.8-Cre mice were crossed with AR^flox/flox^ mice to produce sensory neuron-selective AR-cKO (Nav1.8-Cre::AR^flox/flox^) mice. (b) Mechanical pain thresholds in female WT and AR-cKO mice on day 1 after i.p. administration of DHT (100 mg/kg) were assessed using the up-down method with the von Frey test (n=7-8, two-way ANOVA followed by Sidak’s multiple comparison test, **P=0.0016). (c) Double immunostaining of AR and neuronal markers (NeuN, NF200, CGRP, or IB4-labeled non-peptidergic C-fibers) in the DRG of female WT and AR-cKO mice on day1 after i.p. administration of DHT. Scale bars = 50 μm. (d) Quantitative analysis of the percentage of AR^+^ cells colocalized with neuronal marker^+^ cells in the DRG of female WT and AR-cKO mice (n=6 mice, Mann-Whitney’s U test, **P=0.0022 (NeuN), *P<0.0043 (NF200), **P=0.0022 (CGRP)).

## Discussion

### Sex- and androgen-dependent differences in various pain modalities

In this study, we demonstrated that female mice have lower mechanical pain thresholds compared to male mice, while no significant sex differences were observed in response to heat or chemical pain. GDX male mice also exhibited significantly lower thresholds to mechanical painful stimuli. This was further confirmed by the increased excitation of pain-responsive SDH neurons, which was enhanced under conditions of lower androgen levels in GDX males. These findings suggest that intrinsic androgen levels play a key role in regulating pain sensitivity. In agreement with this, systematic reviews of sex differences in pain perception consistently show that the most notable difference is in the sensitivity to mechanical pain, with females exhibiting lower mechanical pain thresholds compared to males in humans ^4,6^. Although some studies report sex differences in responses to heat, cold, or chemical painful stimuli ^4,6^, such findings remain controversial, and these areas are open to further investigation.

Testosterone has been reported to alleviate muscle pain in both animals and humans, but a variety of cell populations, such as immune cells, are involved in the pathology of chronic pain ^27–29^. Additionally, testosterone levels are altered following tissue injury, leading to pain hypersensitivity. This change correlates with the activation of brain regions by nociceptive stimuli ^30^, and lower testosterone levels have been associated with decreased activity of the descending pain inhibitory system, thereby increasing pain severity ^31^. These lines of evidence demonstrate that systemic exposure to androgens exerts pain-inhibitory effects. However, the cellular target of androgens has been controversial in previous reports and remains to be identified.

### Influence of AR signals on the properties of primary sensory neurons

Several studies have shown that estrogen receptors are also expressed in pain-processing neurons, suggesting that estrogens may contribute to pain regulation as well ^32–35^. In male mice, testosterone is primarily produced by the testes, which also secrete small amounts of other sex hormones, including estrogens ^14^. Since testosterone is converted to estrogens by aromatase ^13,14^, the indirect influence of estrogens should be considered. Given that DHT, a more potent androgen ^14,36^, is not converted to estrogens, our findings on the effects of DHT support the notion that, among sex hormones, androgens appear to be key regulators of mechanical sensitivity.

AR is expressed throughout the pain transmission pathway (e.g., dorsal root ganglia, spinal cord, and brain areas) (**Fig. S6**) ^15,33^, and androgens are involved in regulating pain perception and analgesic effects ^37,38^. In this study, we demonstrated that AR is located in the nuclei of sensory neurons in both normal males and DHT-treated females. Additionally, sensory neuron-selective AR-cKO resulted in a decreased mechanical pain threshold, highlighting the role of AR signaling in regulating pain sensitivity. Since sensory neuron inputs to the SDH enhance the activity of pain-responsive neurons, the increased firing rates of SDH neurons in GDX and AR-cKO males likely represent enhanced pain transmission at the spinal level, correlating with the lower pain thresholds observed in behavioral analyses.

Similar effects of testosterone and DHT administration were observed in female mice, and but these effects were diminished in sensory neuron-selective (Nav1.8-Cre::AR^flox/flox^) AR-cKO females, while they remained unchanged in SDH neuron-selective (Lbx1-Cre::AR^flox/flox^) AR-cKO females. This suggests that the systemic androgens increase mechanical pain thresholds, at least in part, through AR signaling in sensory neurons. Several reports have demonstrated that androgens produce sex differences in various cell types, including immune cells ^39–41^. We assume that androgens alter the physiological properties of these sensory neurons, as previously reported: androgens can directly regulate neural activity^42–44^ and synaptic density ^45–47^.

### Roles of CGRP^+^ neurons in regulating pain sensitivity

Notably, 48% of DRG neurons, including both A- and C-fibers of various sizes, expressed AR in their nuclei (**Fig. 3**). These findings align with a previous report showing nuclear localization of AR in A- and C-fibers of DRG neurons innervating the pelvic regions (L6-S1) ^18^. Approximately 80% of CGRP^+^ neurons expressed AR in normal males as well as in DHT-treated females, consistent with previous findings showing a similar proportion of CGRP^+^ neurons expressing AR ^18^. In AR-cKO males, the highest AR depletion was observed in CGRP^+^ neurons compared to other populations, suggesting that CGRP^+^ neurons expressing AR are critical for the regulation of mechanical pain sensitivity.

Among the various subsets of sensory neurons, CGRP^+^ neurons respond to mechanical pain stimuli ^21^, and CGRP administration induces mechanical hypersensitivity ^48,49^. Interestingly, some reports suggest that female mice are more susceptible to CGRP than males. Administration of small amounts of CGRP produced pain behavior only in females, while larger doses induced pain behaviors in both sexes ^48,49^. Our present findings, along with these reports, indicate that CGRP^+^ neurons are involved in the sex differences in pain sensitivity. This argument is further supported by the fact that CGRP is recognized as a causative molecule for migraines, which are more prevalent in females, and it serves as a target for migraine treatment ^50,51^.

AR is a nuclear receptor that regulates various types of gene expression ^41,52,53^. It is hypothesized that AR may modify a specific set of genes in sensory neurons. Supporting this assumption, androgens have been shown to regulate the expression and/or activity of ion channels, such as transient receptor potential channels ^54^. Additionally, androgens influence G-protein-coupled receptors that directly mediate sensory neuron activity. For example, the expression level of the mu-opioid (MOP) receptor in neurons is regulated by androgens, with AR binding to promoter regions in the trigeminal ganglia. Treatment with DHT increased MOP expression, while blockade of AR signaling decreased it ^33,55,56^. Since these molecules are expressed in CGRP^+^ neurons ^21,56,57^, it is hypothesized that AR signaling is involved in pain perception in CGRP^+^ neurons. Our preliminary transcriptome and gene ontology analyses of AR-cKO mice suggest the possibility that AR signaling may regulate the specific gene expression associated with neuronal activity in CGRP^+^ neurons (unpublished). Identifying pain-related genes regulated by AR may provide insight into the mechanisms underlying sex differences in pain.

### Androgen-dependent plastic regulation of pain sensitivity in both sexes

To explore the potential role of androgens in pain therapeutics, it is crucial to determine whether sex differences in pain sensitivity are attributed to innate factors such as sex-specific genes. Sex-biased anatomical differences are not prominent during the early embryonic stage ^58^; sexual differentiation begins with the surge of testosterone around birth ^59^. Subsequently, testosterone levels rise around postnatal day 30, coinciding with puberty and potentially causing sex differences in several cell types ^59,60^. It is important to determine whether manipulation of androgen levels after sex differences have been established can alter pain sensitivity.

In the present study, GDX performed in males at 4 weeks of age altered pain sensitivity, indicating that the characteristics of sensory neurons may remain flexible regardless of genetic factors. Furthermore, administration of DHT to females at 8 weeks of age or older induced nuclear translocation of AR in the DRG and increased mechanical pain threshold. This suggests that androgens have the potential to alter sensory neuron function even after maturity, thereby influencing pain sensitivity when androgen signaling is manipulated. These findings align with previous reports indicating that treatment with flutamide, an AR antagonist, increased pain severity in adult rats ^37^. The increase in AR expression in the DRG following androgen exposure in females parallels that observed in males, and androgen administration in females can induce a male-like pain response through the action of androgens on primary sensory neurons. Therefore, manipulating these androgenic effects could lead to novel therapeutics aimed at modulating pain sensitivity and tolerance.

### Conclusion

In summary, we have demonstrated for the first time that primary sensory neurons are key targets of androgens in regulating mechanical pain sensitivity. Our findings emphasize not only the strong relationship between androgen levels and mechanical pain thresholds but also that interventions aimed at modulating AR in sensory neurons can effectively control mechanical pain regardless of sex. Further investigations are needed to identify pain-modulating molecules or factors that fluctuate downstream of AR signaling.

## Materials and Methods

### Mice

All animal experiments were approved by the Animal Research Committee of Wakayama Medical University and conducted in accordance with the in-house guidelines for the care and use of laboratory animals at Wakayama Medical University, as well as the Animal Research: Reporting of In Vivo Experiments (ARRIVE) guidelines. Nav1.8 (*Scn10a*)-Cre knock-in mice [B6.129(Cg)-*Scn10a^tm2(cre)Jwo^*/TjpJ; stock #036564] ^23^ purchased from the Jackson Laboratory, and Lbx1-Cre knock-in mice ^25,26^ were crossed with AR^flox/flox^ mice ^24^ or R26-LSL-tdTomato mice ^61^. For the Cre-dependent deletion of the *Ar* gene, located on the X chromosome, in Nav1.8-expressing primary sensory neurons or Lbx1-expressing SDH neurons, floxed-AR sequences were maintained in heterozygous and homozygous genotypes, respectively. Subsequently, 8- to 12-week-old male Nav1.8-Cre::AR^flox/y^ mice, as well as female Nav1.8-Cre::AR^flox/flox^ and Lbx1-Cre::AR^flox/flox^ mice, were used for the experiments. Littermate male AR^flox/y^ and female AR^flox/flox^ mice were used as WT controls. Male and female C57BL/6 mice (4–8 weeks old) were purchased from SLC (Hamamatsu, Japan) and used for experiments at 8–12 weeks of age. All mice were housed in groups of 5–6 in plastic cages under controlled temperature (23–24°C), humidity (60– 70%), and a 12-hour dark/light cycle, with free access to food and water.

### Gonadectomy

For orchiectomy, male mice were anesthetized with isoflurane. After aseptic preparation of the surgical site, a small incision was made in the lower abdomen to exteriorize the testes, vas deferens, and spermatic blood vessels. The blood vessels and vas deferens were cauterized, and the testes were removed either bilaterally (**Fig. 1-3**) or unilaterally (**Fig. 1a, b**). The skin incision was closed with sutures, and the surgical area was sterilized with povidone-iodine. Four weeks post-surgery, the animals were used for experiments ^17^.

### Serum testosterone measurement

Mice were euthanized by decapitation, and trunk blood was collected into separate microtubes (Sansho, Tokyo, Japan). Following the manufacturer’s instructions, the collected blood samples were kept in the tubes for 30 minutes before centrifugation at 4°C for 5 minutes. The upper serum phase was then carefully transferred to a fresh tube. Serum testosterone levels were measured using liquid chromatography-tandem mass spectrometry, which was performed by Asuka Pharmaceutical (Tokyo, Japan).

### Drug administration

TP (Fujifilm Wako, Osaka, Japan) and DHT (Tokyo Chemical Industry, Tokyo, Japan) were dissolved in a dimethyl sulphoxide/sesame oil solution at a ratio of 1:9. These drugs were administered intraperitoneally (i.p.) at a volume of 0.05 ml/10 g body weight to awake mice.

### von Frey test

Mechanical pain threshold was determined using the von Frey test, as previously described ^62^. Briefly, mice were individually placed on a metal mesh grid floor (5 × 5 mm) and covered with an acrylic box. After a 2-to 3-hour adaptation period, calibrated von Frey filaments (Neuroscience, Tokyo, Japan) were applied to the middle of the plantar surface of the hind paw through the mesh floor. The filament set used in this study consisted of nine calibrated von Frey filaments: 0.02, 0.04, 0.07, 0.16, 0.4, 0.6, 1.0, 1.4, and 2.0 g. Using the up-down method, testing began with the application of 0.4 g filament. Quick withdrawal, shaking, biting, or licking of the stimulated paw was considered a positive paw-withdrawal response. If no withdrawal response occurred, the next stronger stimulus was applied. Conversely, the next weaker stimulus was selected following a paw withdrawal, in accordance with Chaplan’s procedure ^63^. Once the response threshold was crossed (two responses were straddling the threshold), the 50 % paw-withdrawal threshold was calculated based on these responses.

### Hargreaves test

Thermal pain sensitivity was assessed using the Hargreaves test, as previously described ^64^. Mice were individually placed on a glass sheet and covered with a clear acrylic box. After a 2-to 3-hour adaptation period, a radiant heat source (IITC 390 Plantar Test Analgesia Meter, Neuroscience) was applied to the plantar surfaces of both hind paws. Withdrawal latencies were calculated as the mean latency of three stimulations. A cut-off latency of 15 seconds was set to avoid tissue damage.

### Formalin test

Chemical pain intensity was assessed using the formalin test. Mice were individually placed in a clear acrylic box and allowed to acclimate to the environment for 60 minutes. Following an intraplantar injection of 2% formalin solution (20 μl) into the left hind paw of awake mice, their behavior was videotaped for 60 minutes. Pain-related behaviors, such as licking, biting, lifting, or flinching of the injected paw, were measured at 5-minutes intervals by reviewing the videotaped footage. All behavioral assessments were conducted in a blinded manner.

### Rotarod test

A rotarod apparatus (Panlab, Barcelona, Spain) was used to assess the motor function. A few days prior to testing, the mice underwent pretraining to habituate to the rod rotating at different speeds (5, 10, and 15 rpm). During the test, mice were placed on the rod facing the direction opposite to the rotation and allowed to ambulate until they fell off the rod or the maximum observation time (180 seconds) was reached. For all experiments, the latency to fall and the rotational velocity at which the mice fell from the rod were recorded. After several days of training, the mice rested for one day before being tested on the rod at various speeds (5, 10, and 15 rpm). The mean time spent on the rotarod at each velocity was measured.

### In vivo extracellular recordings

*In vivo* extracellular recordings were performed as described in previous reports ^61,65^. Briefly, mice were anesthetized via i.p. administration of urethane (1.2–1.5 g/kg). The procedure involved the removal of thoracolumbar vertebral arches, exposing the Th11–L4 vertebrae, after which the animals were secured in a stereotaxic frame. The dura mater was excised, and the arachnoid membrane was incised to allow for the insertion of tungsten microelectrodes. The spinal cord surface was continuously bathed with Krebs solution, equilibrated with 95% O_2_ and 5% CO_2_ (flow rate: 10–15 ml/min), maintained at a temperature of 37 ± 1 ◦C. The solution contained NaCl, KCl, CaCl_2_, MgCl_2_, NaH_2_PO_4_, glucose, and NaHCO_3_. For extracellular single-unit recordings from deep dorsal horn neurons (specifically lamina III–IV), recordings were obtained at depths ranging from 180 to 400 μm below the surface. The unit signals were amplified (EX1, Dagan Corporation, Minneapolis, MN, USA), digitized (Digidata 1400A; Molecular Devices, Union City, CA, USA), and analyzed using Clampfit software (version 10.2; Molecular Devices). Tactile stimuli were applied using excised cotton, and noxious pinch stimuli were delivered using forceps to identify responsive neural regions. Mechanical stimulation involved bending the skin with von Frey filament, applying forces of 0.16, 0.4, and 1.0 g. A 5-second stimulus was administered to the most responsive sites on each hind limb. For lidocaine testing, a 0.5% solution (AstraZeneca, Osaka, Japan) was applied to the stimulation points.

### Immunohistochemistry

The lumbar (L4–5) DRG and the spinal cord were harvested from euthanized mice following transcardiac perfusion with phosphate-buffered saline (PBS) and fixed in 4% paraformaldehyde/phosphate buffer solution. The specimens were post-fixed in 4% paraformaldehyde and cryoprotected in 30% sucrose/PBS solution at 4°C overnight. After embedding in a freezing compound (Sakura, Tokyo, Japan), frozen tissues were longitudinally sectioned at 12 μm thickness (DRG) or 30 μm thickness (spinal cord) using a cryostat (Leica Microsystems, Wetzlar, Germany). DRG sections were mounted on glass slides, while spinal cord sections were floated in PBS. The sections were treated with PBS containing 0.1 % Triton X-100 (PBST) for 1 hour and then blocked with 5% donkey serum at room temperature (15–25°C) for 2 hours. They were then incubated overnight at 4°C with primary antibodies targeting the AR (rabbit polyclonal, 1:100; Abcam, Cambridge, UK), NeuN (mouse monoclonal, 1:500; Millipore, Billerica, MA, USA), NF200 (mouse monoclonal, 1:500; Millipore), and CGRP (goat polyclonal, 1:2000; Abcam). The sections were rinsed in PBST and incubated with fluorescence-conjugated secondary antibodies (1:200; Thermo Fisher Scientific, Waltham, MA, USA) or isolectin-B4 (IB4, Thermo Fisher Scientific) at room temperature for 2 hours. Finally, the sections were washed with PBS, mounted on glass slides, and covered with coverslips using Immunoselect Antifading Mounting Medium DAPI (Dianova, Hamburg, Germany). Fluorescent images were acquired using a confocal laser scanning microscope (Olympus, Tokyo, Japan). The number of AR^+^ cells and the colocalization of AR with neuronal markers in the DRG were quantified across all sections using FLUOVIEW software ^17^.

### Gene expression analysis

Expression profiling of Nav1.8 (*Scn10a*) and AR (*Ar*) genes across a diverse range of normal tissues, organs, and cell lines in mice was visualized using BioGPS (http://biogps.org/).

### Statistical analysis

All data are presented as the mean ± standard error of the mean (SEM). Shapiro-Wilk’s test and Levene’s test were used to confirm the homogeneity of variance. To compare differences between the two groups, two-tailed Student’s t-test (**Fig. 1c**, **Fig. 4c**), Welch’s t-test (**Fig. 4b**, **Fig. 6b, Fig. S3**), or Mann-Whitney U test (**Fig. 1d**, **Fig. 2a-d**, **Fig. 4d,f**, **Fig. 5a,b**, **Fig. 6d**, **Fig. 7d, Fig. S4**) was used. For comparisons among three or more groups, one-way analysis of variance (ANOVA) followed by Dunnett’s multiple comparison test (**Fig. 1a**, **Fig. 6a**) or Kruskal-Wallis test followed by Dunn’s multiple comparison test (**Fig. 1b**, **Fig. 3b**) was employed. To compare differences among four groups with two factors, two-way ANOVA followed by Sidak’s multiple comparison test (**Fig. 7b, Fig. S5d**) was used. Statistical analyses were performed using GraphPad Prism software (GraphPad Software, Boston, MA, USA), and statistical significance was set at P < 0.05.

## Acknowledgments

We are grateful to Dr. Gen Yamada (Wakayama Medical University, Wakayama, Japan), Dr. Carmen Birchmeier (Max Delbrück Center for Molecular Medicine, Berlin, Germany), and Dr. Kazuhiko Nishida (Kansai Medical University, Osaka, Japan) for providing mouse lines. We also thank Ms. Mayumi Shibutani for technical assistance and Editage (https://www.editage.com) for English language editing.

## Author contributions

Conceptualization, S.H., K.S., and N.K.; Methodology, F.S., D.U., K.S., and N.K.; Validation, D.U., S.H., K.S., and N.K.; Formal Analysis, D.U., S.H., K.S., and N.K.; Investigation, F.S., D.U., Y.F., Y.Hi., Y.Ha., and N.K.; Resources, D.U., K.S., and N.K.; Data Curation, F.S., D.U., Y.F., Y.Hi., Y.Ha., S.K., H.N., S.H., K.S., and N.K.; Writing-Original Draft, N.K.; Writing-Review & Editing, F.S., D.U., Y.F., Y.Hi., Y.Ha., S.K., H.N., S.H., K.S., and N.K.; Visualization, N.K.; Supervision, N.K.; Project Administration, N.K.; Funding Acquisition, D.U., Y.F., K.S., and N.K.

## Data and materials availability

All data needed to evaluate the conclusions in the paper are present in the paper and/or the Supplementary Materials.

