## Supplementary figures for "Androgen receptors expressed in the primary sensory neurons regulate mechanical pain sensitivity"

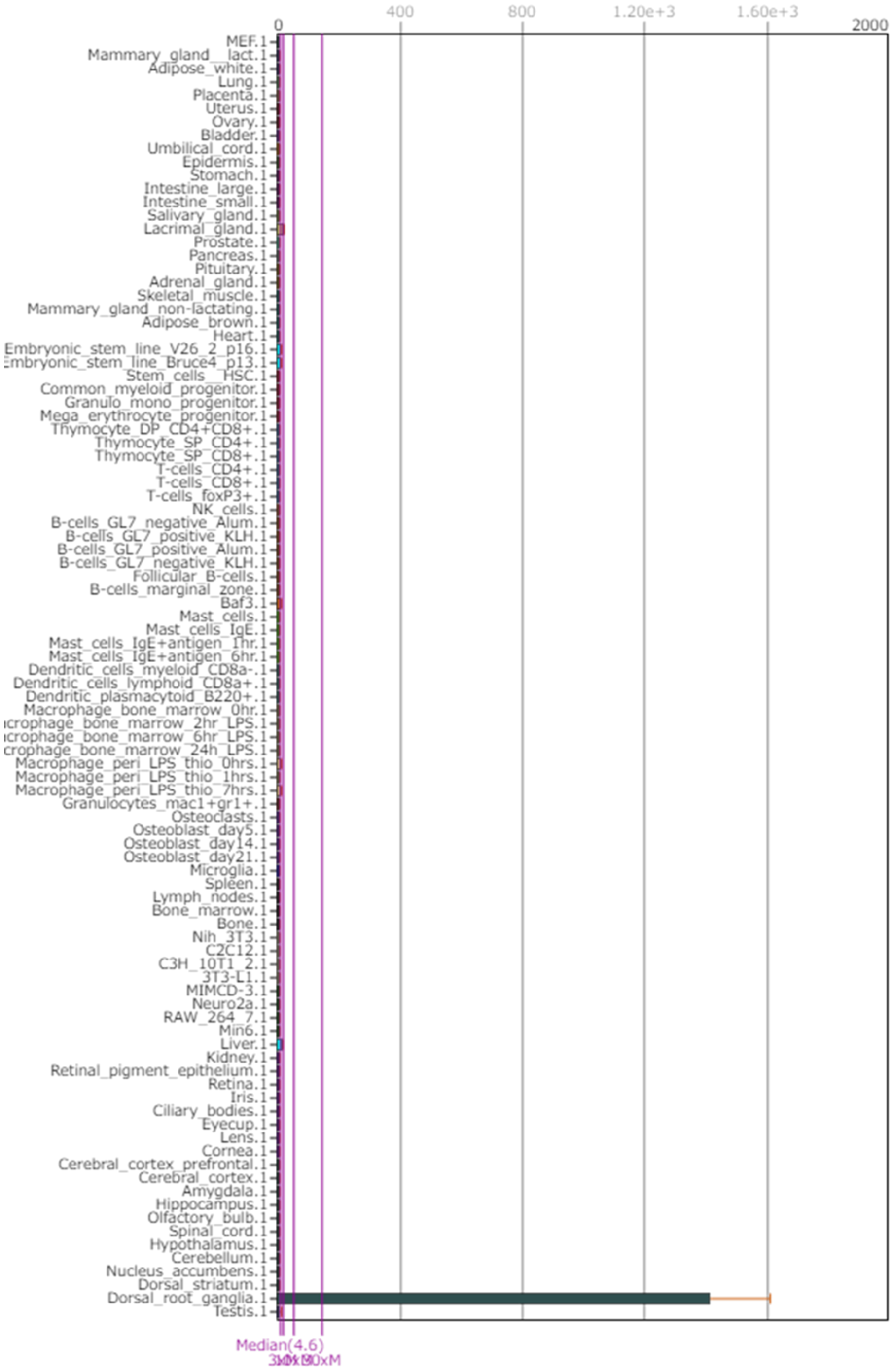


**Fig. S1. Expression profiling of *Scn10a* gene in mice.** *Scn10a* gene expression across a diverse array of normal tissues, organs, and cell lines in mice was visualized using BioGPS. Dataset: GeneAtlas MOE430, gcrma. Probeset: 1450266_at.

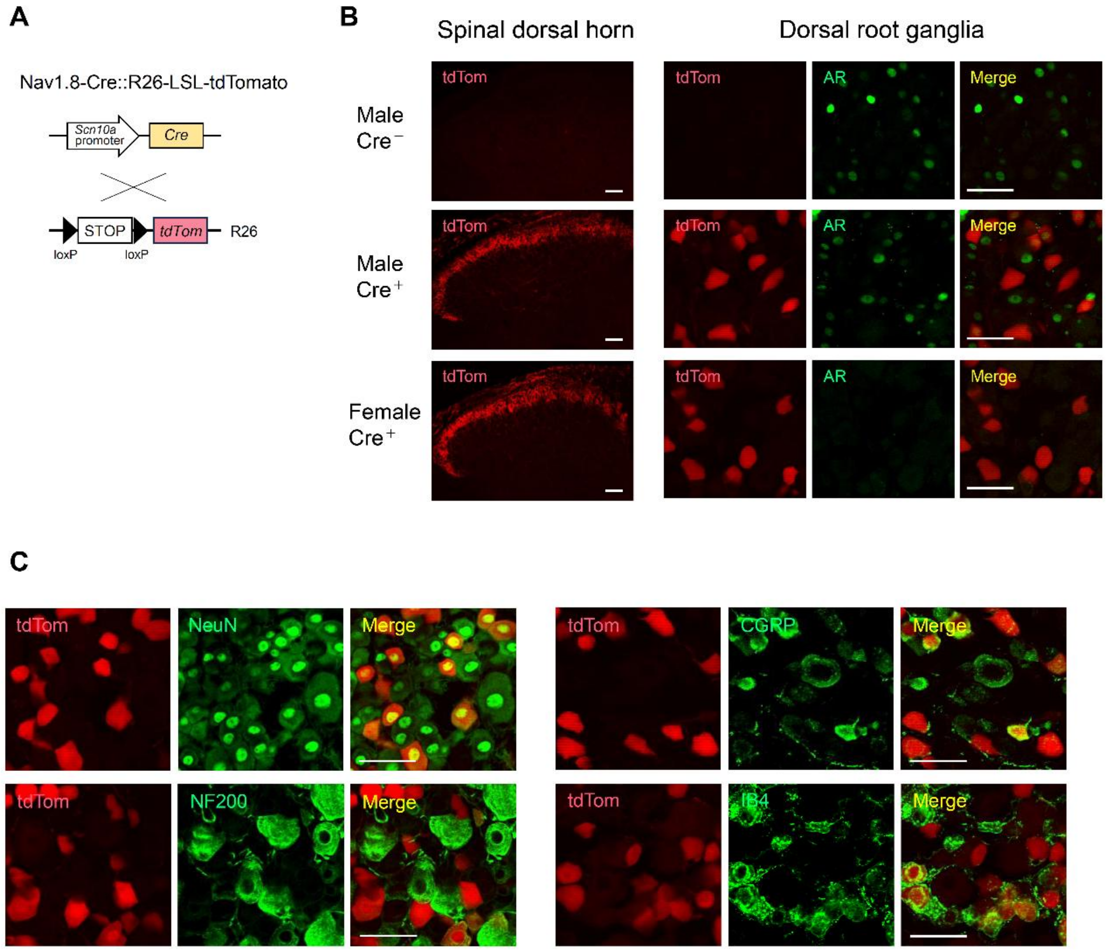


**Fig. S2. Expression of Cre recombinase in the sensory neurons in males and females.** (a) Nav1.8-Cre mice were crossed with R26-LSL-tdTomato mice to visualize Cre-expressed sensory neurons. (b) Expression of tdTomato (tdTom) in the spinal dorsal horn (SDH) and dorsal root ganglia (DRG) in male Cre$^-$, male Cre$^+$, and female Cre$^+$ mice was visualized by immunohistochemistry. (c) Expression of tdTomato and neuronal markers (NeuN, NF200, CGRP, or IB4-labeled non-peptidergic C-fibers) in the DRG of Male Cre$^+$ mice. Scale bars = 50 μm (b,c).

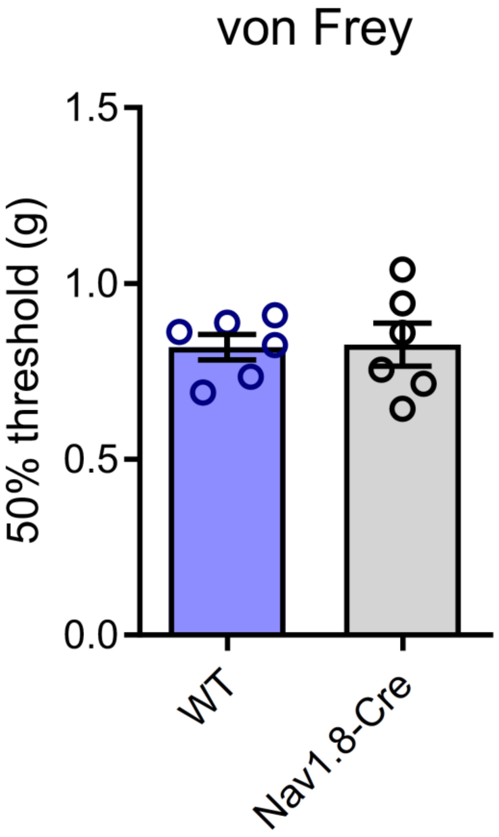


**Fig. S3. Pain sensitivity in male Nav1.8-Cre mice.** Mechanical pain thresholds in male wild-type (WT) and Nav1.8-Cre mice were assessed using the up-down method with the von Frey test (n=6, Welch's t-test).

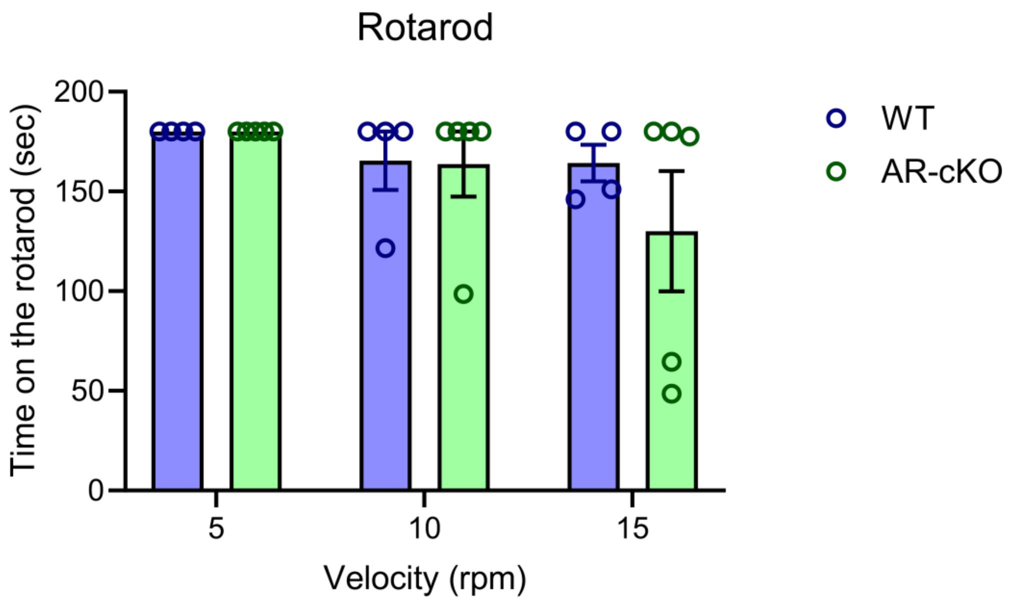


**Fig. S4. Motor function in male sensory neuron-selective androgen receptor conditional knockout (AR-cKO) mice.** The mean time spent on the rotarod at each velocity in male WT and AR-cKO mice was measured using the rotarod test (n=4-5 mice, Mann-Whitney's U test).

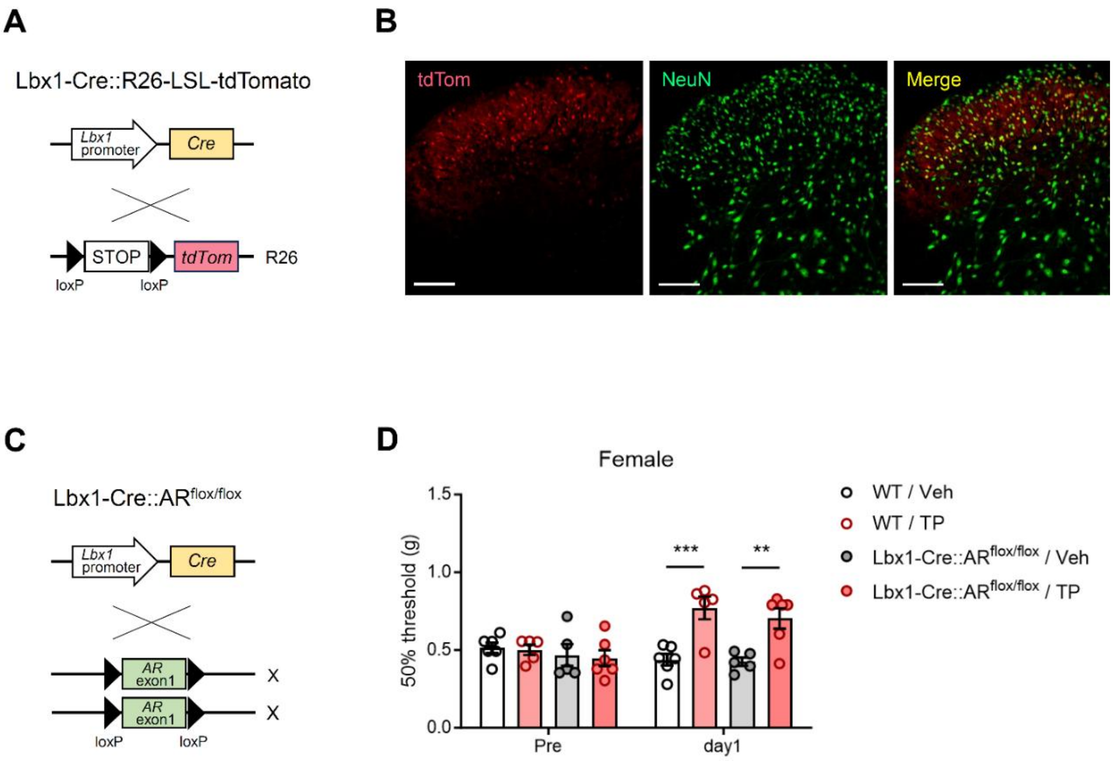


**Fig. S5. Contribution of AR in SDH neurons to regulating mechanical pain sensitivity in females.** (a) Lbx1-Cre mice were crossed with R26-LSL-tdTomato mice to visualize Cre-expressed SDH neurons. (b) Expression of tdTomato (tdTom) in the SDH of female Cre$^{+}$ mice were visualized by IHC. Scale bars = 100 μm. (c) Lbx1-Cre mice were crossed with AR$^{flox/flox}$ mice to generate SDH neuron-selective AR-cKO (Lbx1-Cre::AR$^{flox/flox}$) mice. (d) Mechanical pain thresholds in female WT and AR-cKO mice were assessed on day 1 after i.p. administration of dihydrotestosterone (DHT; 100 mg/kg) using the up-down method with the von Frey test (n=5–6, two-way ANOVA followed by Sidak's multiple comparison test, ***P<0.001, **P=0.0036).

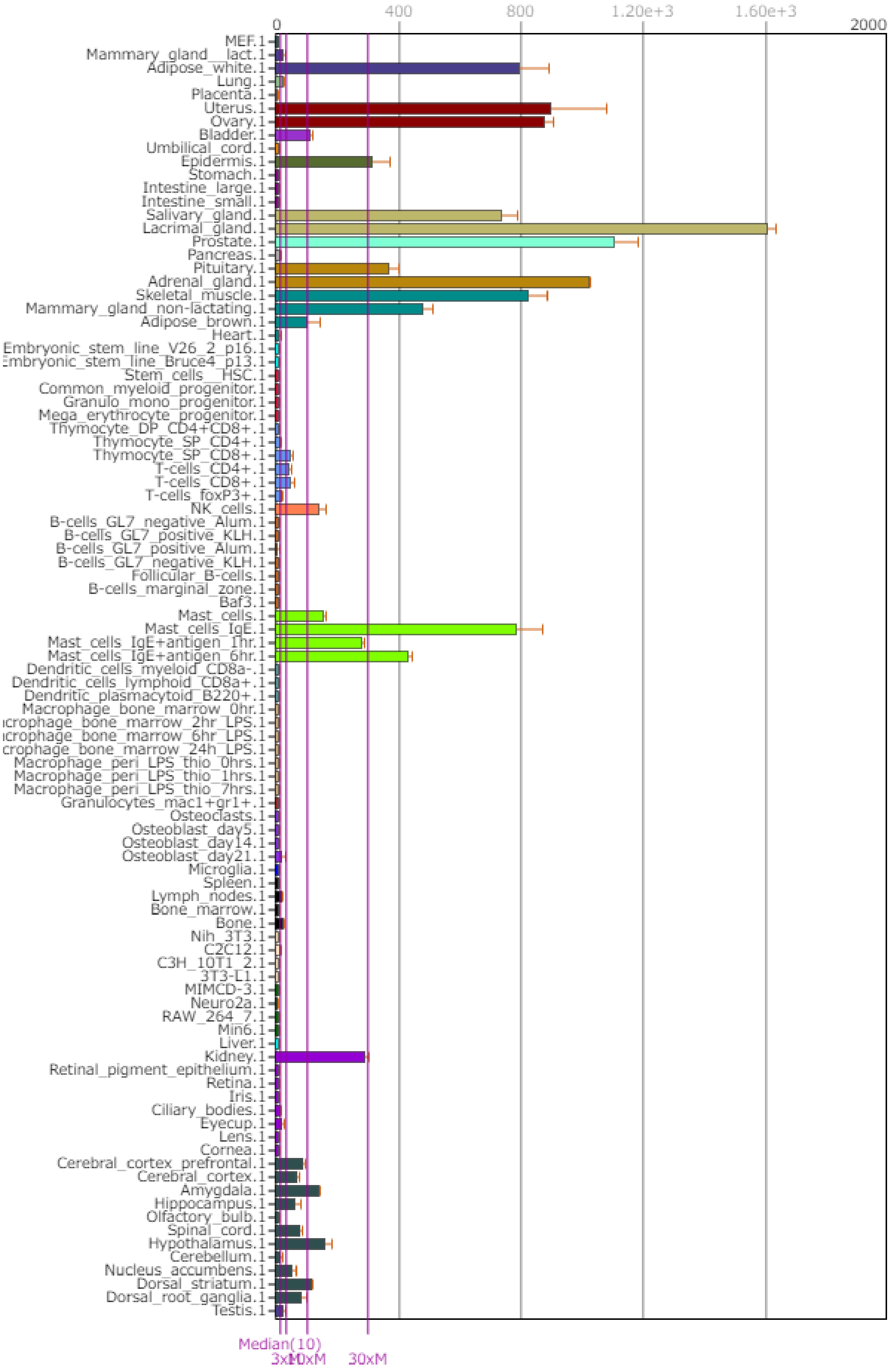


**Fig. S6. Expression profiling of Ar gene in mice.** *Ar* gene expression across a diverse array of normal tissues, organs, and cell lines in mice was visualized using BioGPS. Dataset: GeneAtlas MOE430, gcrma. Probeset: 1437064_at.
